# Extracellular sodium regulates fibroblast growth factor 23 (FGF23) formation

**DOI:** 10.1101/2023.06.23.546282

**Authors:** Zsuzsa Radvanyi, Eun Jin Yoo, Palanivel Kandasamy, Adrian Salas-Bastos, Sophie Monnerat, Julie Refardt, Mirjam Christ-Crain, Himeka Hayashi, Yasuhiko Kondo, Jonathan Jantsch, Isabel Rubio-Aliaga, Lukas Sommer, Carsten A. Wagner, Matthias A. Hediger, Hyug Moo Kwon, Johannes Loffing, Ganesh Pathare

## Abstract

Fibroblast growth factor-23 (FGF23) is a bone-derived hormone that has recently received much attention due to its association with the progression of chronic kidney disease, cardiovascular disease, and associated mortality. Extracellular sodium concentration ([Na^+^]) plays a significant role in bone metabolism. Hyponatremia (low serum [Na^+^]) has recently been shown to be independently associated with FGF23 levels in patients with chronic systolic heart failure. However, nothing is known about the direct impact of [Na^+^] on FGF23 production. Here, we show that an elevated [Na^+^] (+20 mM) suppressed FGF23 formation, whereas low [Na^+^] (−20 mM) increased FGF23 synthesis in the osteoblast-like cell line UMR-106. Similar bidirectional changes in FGF23 abundance were observed when osmolality was altered by mannitol but not by urea, suggesting a role of tonicity in FGF23 formation. Moreover, these changes in FGF23 were inversely proportional to the expression of NFAT5 (nuclear factor of activated T cells-5), a transcription factor responsible for tonicity-mediated cellular adaptations. On the other hand, arginine vasopressin (AVP), which is often responsible for hyponatremia, did not affect FGF23 production. Next, comprehensive and unbiased RNA-seq analysis of UMR-106 cells exposed to low vs. high [Na^+^] revealed several novel genes involved in cellular adaptation to altered tonicity. Additional analysis of cells with Crisp-Cas9 mediated NFAT5 deletion indicated that NFAT5 controls numerous genes associated with FGF23 synthesis, thereby confirming its role in [Na^+^]-mediated FGF23 regulation. In line with these in vitro observations, we found that human hyponatremia patients have higher FGF23 levels. Our results suggest that [Na^+^] is a critical regulator of FGF23 synthesis.

**SIGNIFICANCE STATEMENT:** Fibroblast growth factor 23 (FGF23) is a bone-derived hormone that controls phosphate and vitamin D metabolism. Excess FGF23 is postulated to cause left ventricular hypertrophy, while FGF23 deficiency reduces life span and mimics age-related diseases in mice. FGF23 is also a potential biomarker for chronic kidney disease and cardiovascular disorders, but its role in disease progression is unclear. Therefore, it is important to explore the regulation of FGF23 production, which is incompletely understood. Our paper identifies extracellular-sodium-NFAT5 signaling as a key regulator of FGF23 formation.

## INTRODUCTION

Fibroblast growth factor-23 (FGF23) was discovered as markedly elevated ‘phosphatonin’ in patients with autosomal dominant hypophosphataemic rickets (ADHR)^1^. High levels of FGF23 were also found in X-linked hypophosphatemia (XLH), which is caused by inactivating mutations in the *Phex* gene^2, 3^. Eventually, it was found that FGF23 regulates phosphate and vitamin D homeostasis by inhibiting renal sodium phosphate cotransporters and suppressing the process of vitamin D biosynthesis, respectively^4^. As a result, disruption of the FGF23 endocrine axis plays a key role in the pathophysiology of renal and bone disorders as well as aging^4^. In recent years, FGF23 has gained significant interest due to its strong association with poor prognosis in chronic kidney disease (CKD) and cardiovascular disease (reviewed in^5^). In mice, high FGF23 induces left ventricular hypertrophy^6^. FGF23 is mainly produced by osteoblasts and osteocytes. Upon secretion, it undergoes cleavage, leading to the presence of both intact (iFGF23) and C-terminal fragments (cFGF23) in the circulation^7^. Dietary phosphate^8, 9^, vitamin D^8^, insulin^10^, volume-regulation^11, 12^, aldosterone^13, 14^, iron status^15^ and inflammation^16^ have been identified as endogenous regulators of FGF23 in bone. However, nothing is known regarding the role of extracellular sodium ion concentration ([Na+]) or osmolality in regulating FGF23 synthesis.

Both hyponatremia (serum [Na^+^] <135 mM) and hypernatremia (serum [Na^+^] >145 mM) can affect bone remodeling. Several studies have provided compelling evidence linking hyponatremia to bone loss, osteoporosis, and heightened bone fragility^17–22^. Most hyponatremia cases are due to the syndrome of inappropriate antidiuretic hormone secretion (SIAD), which is characterized by high arginine vasopressin (AVP) levels (reviewed in^23^). On a mechanistic level, both high AVP^24, 25^ and low [Na+] levels^17, 26^ are postulated to promote osteoclastogenesis and inhibit osteoblastogenesis, leading to bone loss in hyponatremia. Correction of hyponatremia in hospitalized patients has a positive impact on osteoblast function^27, 28^. On the other hand, high [Na^+^] may also enhance bone loss through an increase in osteoclastic resorption^29^. When cells were exposed to high [Na^+^], the expression of the RANKL decoy receptor osteoprotegerin (*Opg*) increased in both osteoclast-precursor cells and osteoblasts^30^, suggesting a direct effect of elevated [Na^+^] on bone.

The intracellular milieu is almost immediately affected by changes in [Na^+^] or osmolality. Within cells, osmoregulation is mainly governed by the tonicity-responsive transcription factor tonicity-responsive enhancer-binding protein (TonEBP), also called the nuclear factor of activated T cells 5 (NFAT5)^31^. Hypertonic conditions induce the upregulation of NFAT5, leading to the transcription of numerous NFAT5 target genes associated with adapting to high [Na^+^]^31^. Nevertheless, NFAT5 remains active even under isotonic conditions and can be either upregulated or downregulated in response to changes in tonicity^32^. A recent genome-wide association study (GWAS) on plasma [Na^+^] concentration identified genetic variants in NFAT5^33^. This suggests that NFAT5 may participate in the regulation of systemic [Na^+^] / water balance.

A recent study has shown that hyponatremia is independently associated with FGF23 levels in patients with chronic systolic heart failure^34^. Given the important role of [Na^+^] in osteoblast functions, we hypothesized that the altered [Na^+^] levels may regulate the production of FGF23 through NFAT5. By manipulating culture media, we studied whether altered [Na^+^] mediated tonicity or osmolality, or both were responsible for FGF23 secretion by osteoblasts. Additionally, we studied the regulation of FGF23 by high AVP, which is often observed in hyponatremic patients.

## RESULTS

### Elevated [Na^+^] mediated hypertonicity suppresses FGF23 formation

NaCl, as an impermeable solute, creates a hypertonic environment when its extracellular levels increase. NFAT5 is the key transcription factor involved in the adaptation to hypertonicity^31^. As demonstrated in Fig. 1A/B/C, +NaCl (20 mM) and mannitol (40 mM), significantly increased NFAT5 protein and mRNA expression in the rat osteoblast-like cell line UMR-106. However, 40 mM urea did not affect NFAT5 protein/ mRNA expression since it elevates osmolality rather than tonicity via cell membrane permeability. The osmolality of normal DMEM cell culture media was 300.7 ± 2.0 mOsm/Kg. There was an almost similar increase in osmolality after the addition of +NaCl, mannitol, and urea (Table. S1). Importantly, the total FGF23 (cFGF23) in the cell supernatant was markedly suppressed by +NaCl and mannitol, but not by urea (Fig. 1D). Similarly, in Fig. 1E, Fgf23 mRNA in cells was also significantly suppressed by +NaCl and mannitol, but not by urea. iFGF23 was not detected due to its very low levels in the cell supernatant (not shown). Consistent with previous reports^30^, we confirmed that +NaCl concentration up to ∼80 mM does not affect cell survival (Fig. 1F). The +NaCl effect on both *Fgf23* and *Nfat5* mRNA was concentration-dependent, with as little as 10 mM +NaCl significantly suppressing *Fgf23* mRNA (Fig. 1G). Time-course experiments showed that *Nfat5* mRNA peaked at 8 h of high-NaCl treatment, while the nadir of *Fgf23* mRNA levels was observed at 24 h (Fig. 1H).

**Fig. 1.**
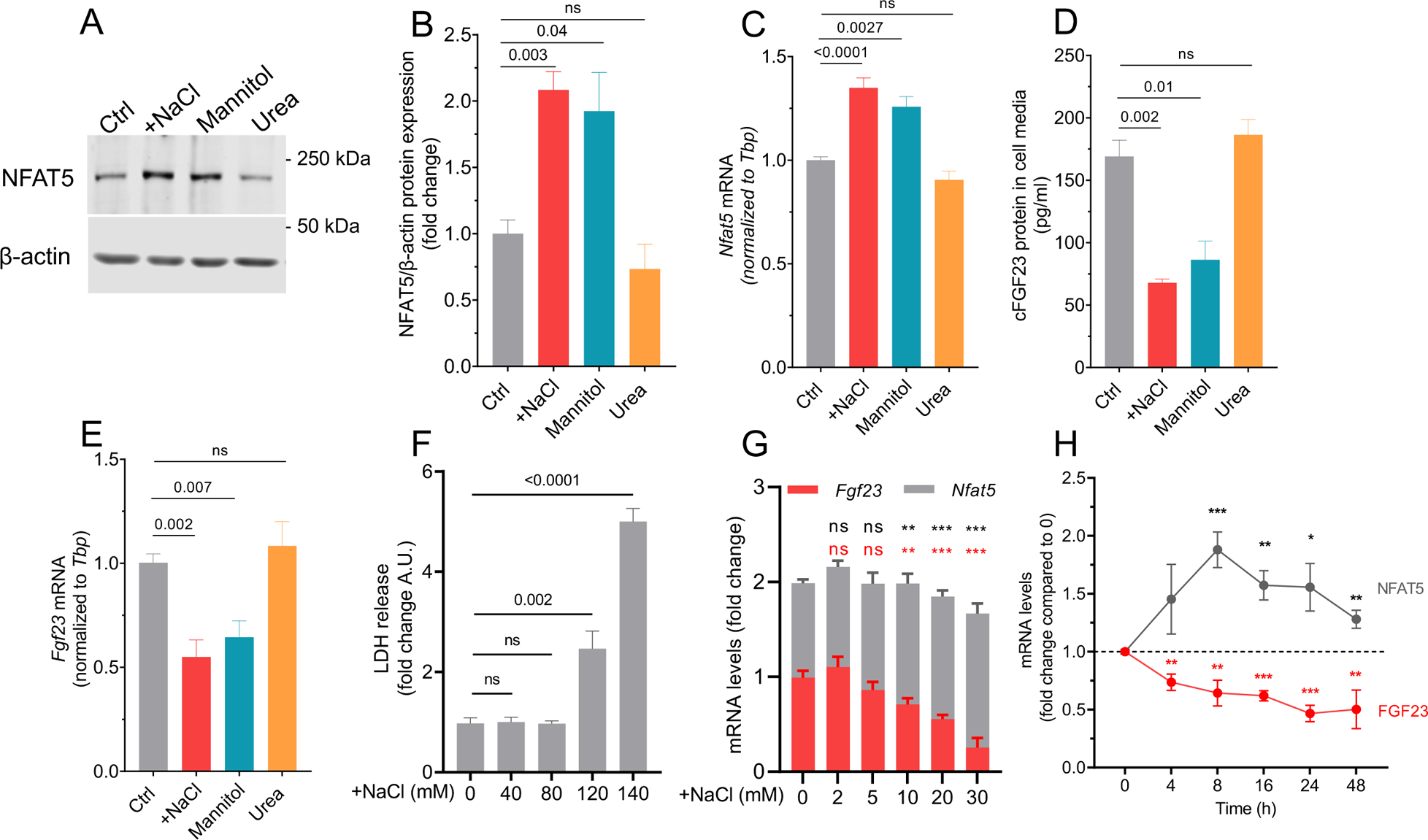
High NaCl suppresses FGF23 formation in UMR-106 cells. **A**) Representative immunoblots of NFAT5 and β-actin upon addition of 20 mM NaCl, 40 mM mannitol, and 40 mM urea treatment for 16 h. B) Quantification of NFAT5 protein normalized with β-actin (n=3). C) Fold change *Nfat5* mRNA levels; D) cFGF23 levels in cell media; E) Fold change *Fgf23* mRNA levels, after treating 20 mM NaCl, 40 mM mannitol, and 40 mM urea for 24 h (n=3-4). F) Lactate dehydrogenase (LDH) release in the cells following 0, 40, 80, 120, and 140 mM +NaCl treatment for 24 h (n=3). G) Fold change *Fgf23* and *Nfat5* mRNA levels following 0, 2, 5, 10, 20, and 30 mM +NaCl treatment for 24 h (n=4). The black and red stars indicate a statistically significant difference between *Nfat5* and *Fgf23* mRNA, respectively, when compared to 0 mM +NaCl. H) Fold change *Fgf23* and *Nfat5* mRNA levels after 20 mM +NaCl treatment over 0-48 h (n=3). The black and red stars indicate a statistically significant difference between *Nfat5* and *Fgf23* mRNA, respectively, when compared to 0 h (control). All the values are expressed in arithmetic means ± SEM. In cases where the *p* value is not mentioned, the following applies: ns (not significant) *p* > 0.05, *\*p* ≤ 0.05, \*\**p* < 0.01, and \*\*\**p* < 0.001.

### Hypotonicity increases FGF23 production

We explored the possibility that hyponatremia might have a direct effect on FGF23 formation. To mimic hyponatremia, cells were cultured in low [Na^+^] media. To generate cell culture media with low [Na^+^] concentrations, custom-made NaCl-free media was reconstituted with specific amounts of NaCl to obtain media in which [Na^+^] were −5, −10, −15, and −20 mM lower than in control media. As shown in Fig. 2A, we found that reducing [Na^+^] in the culture media increased *Fgf23* mRNA levels in a dose-dependent manner. To understand the underlying mechanisms by which low [Na^+^] stimulates FGF23 formation, we manipulated the low [Na^+^] media (−20 mM) with and without correction of osmolality by the addition of 40 mM mannitol or 40 mM urea. As shown in Table S2, the osmolality of low [Na^+^] medium was 260.7 ± 1.5 mOsm/kg. There was an almost similar correction in osmolality after the addition of mannitol and urea (∼302 mOsm/Kg). Interestingly, the addition of mannitol to the low [Na^+^] medium reversed the increase in FGF23; however, the addition of urea did not affect FGF23 formation. This effect was observed at both mRNA and protein levels measured from cells and cell supernatants, respectively (Fig 2B/C). These findings suggested that low tonicity rather than low osmolality was responsible for the increase in FGF23 upon reducing [Na^+^] concentrations. Expectedly, upon reducing [Na^+^] concentration, *Nfat5* mRNA was significantly lower, the effect, again, was reversed by mannitol but not urea (Fig. 2D).

**Fig. 2.**
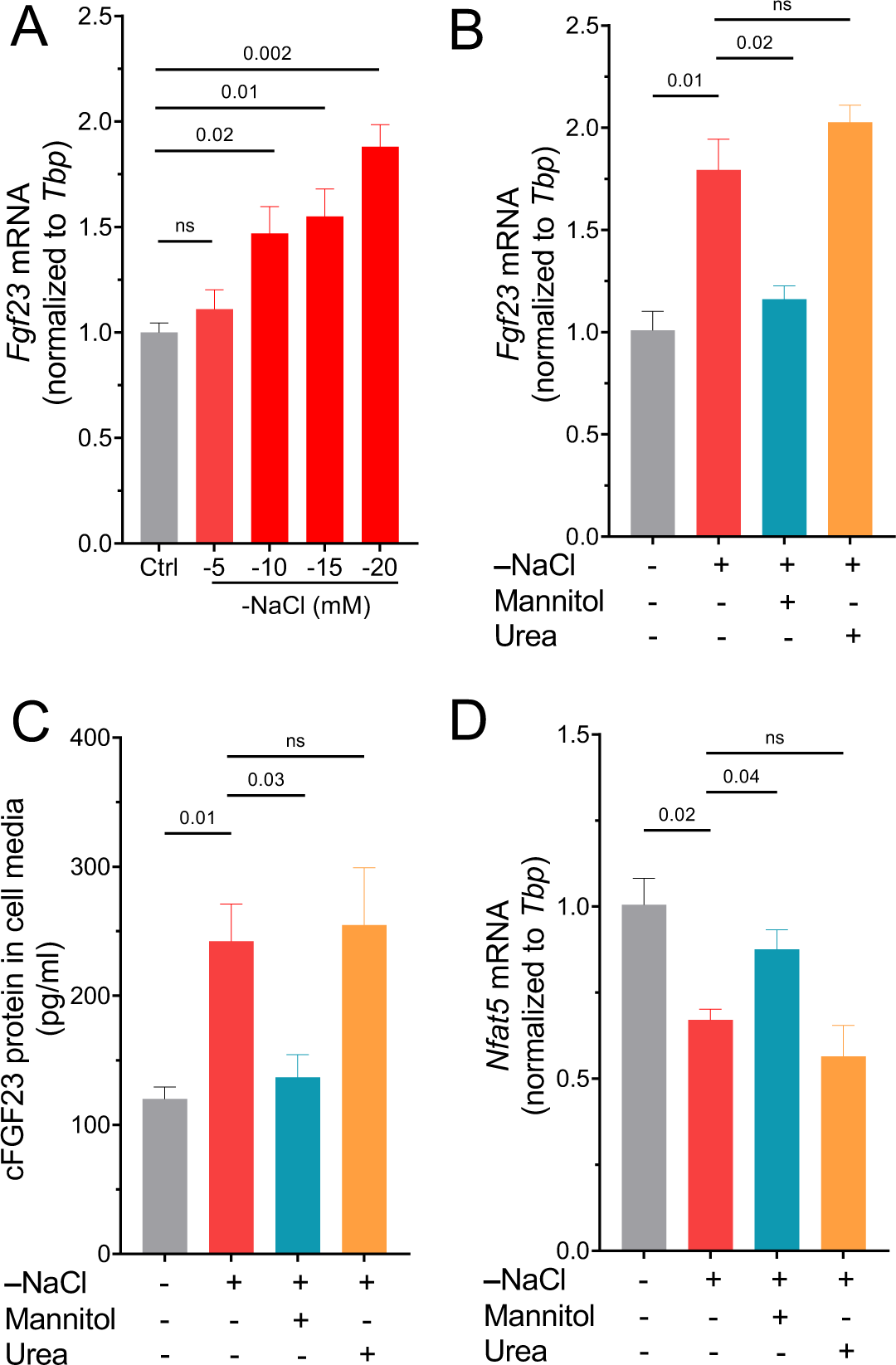
Hypotonicity elevates FGF23 formation in UMR-106 cells. A) Fold change *Fgf23* mRNA levels upon treating NaCl-deficient culture media by −5, −10, −15, and −20 mM NaCl for 24 h (n=3). B) Fold change *Fgf23* mRNA; C) cFGF23 levels in media; D) *Nfat5* mRNA, after treating cells with −20 mM NaCl-deficient culture media. The osmolality was corrected by adding mannitol (40 mM) or urea (40 mM) for 24 h (n=3). All the values are expressed in arithmetic means ± SEM. ns (not significant) *p* > 0.05

### No evidence for FGF23 regulation by AVP

AVP exerts major pathophysiological actions through three distinct receptor isoforms designated V_1a_, V_1b_, and V_2_. Both V_1a_ and V_2_ receptors have been found in murine osteoblasts and osteoclasts, where they may take part in bone remodelling^24, 25^. To this end, we checked the expression pattern of AVP receptors in the rat osteoblast-like UMR-106 cell line. Endpoint PCR was performed on RNA isolated from rat liver (positive control for V_1a_ receptor), kidney (positive control for V_2_ receptor), and UMR-106 cells. As shown in Fig 3A, V_1a_ receptors were very abundant in the liver, while also expressed in kidney tissue as well as UMR-106 cells. However, V_2_ receptors were mostly detected in kidney tissue with only slight expression in liver and UMR-106 cells. To further confirm that V_1a_ is the major AVP receptor expressed in UMR-106 cells, qRT-PCR was performed. As shown in Fig 3B, V_1a_ mRNA levels were ∼7 times higher than V_2_ mRNA in UMR-106 cells. As expected, the pituitary specific-V_1b_ receptor^35^ was undetectable in UMR-106 cells. Next, we treated UMR-106 cells with AVP (1-1000 nM) for 24 hours (Fig. 3C). However, this did not result in any significant change in *Fgf23* mRNA expression. Moreover, AVP (100 nM) applied for various time points between 3-24 h also did not affect *Fgf23* mRNA levels (Fig. 3D). We found that the main AVP receptor expressed in UMR-106 cells is V_1a_; however, AVP binds more strongly to V_2_ than V_1a_ receptors in rats (Kd= 1.7 nm for the V_1a_; Kd= 0.4 nm for the V2)^35^. Therefore, next, we explored the effect of specific V_1a_ agonist ([Phe^2^]OVT, [Phe^2^,Orn^8^]vasotocin)^36^ and V_2_ agonist (ddAVP) on *Fgf23* synthesis. However, both V_1a_ agonist and V_2_ agonist, at low and high concentrations, did not affect *Fgf23* production significantly, when treated for a duration of 24h (Fig. 3E/F). Consistent with these *in-vitro* data, the lack of V_1a_ receptors in mice had no effect on FGF23 levels in the serum (Fig. 3G). Overall, based on these findings, we concluded that although V_1a_ receptors are adequately expressed in UMR-106 cells, AVP does not modulate FGF23 formation.

**Fig. 3.**
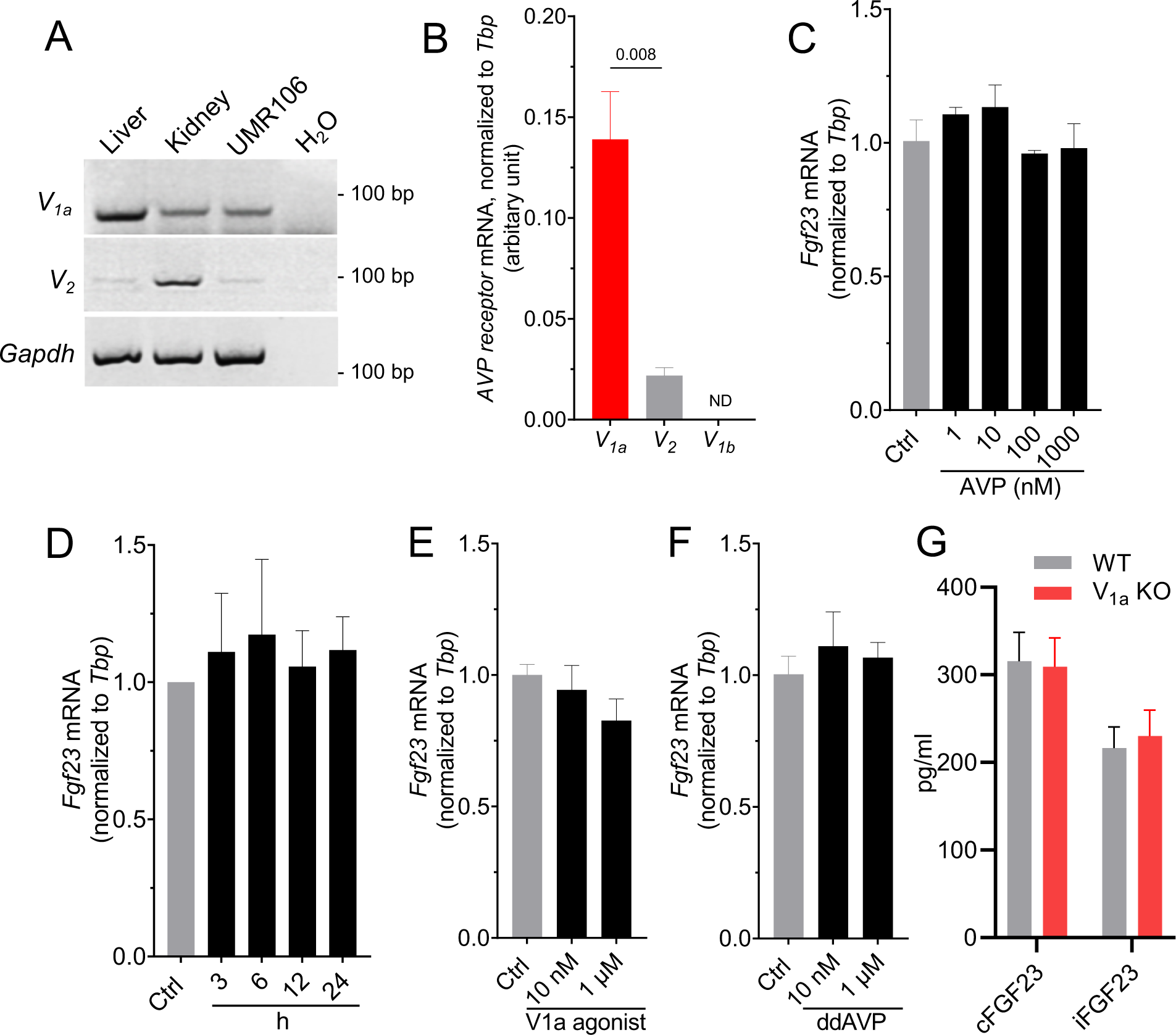
No evidence for FGF23 regulation by AVP. A) Agarose gel electrophoresis for detection of rat V_1a_ and V_2_ receptor mRNA in rat liver, kidney, and UMR-106 cells. The cDNA (2 μg) obtained by RT was amplified for 34 cycles by PCR. B) V_1a_, V_2,_ and V_1b_ mRNA expressions quantified by qRT-PCR in UMR-106 cells (n=3). ND: not detected. C) Fold change *Fgf23* mRNA abundance relative to *Tbp* in UMR-106 cells treated without (Ctrl) or with the indicated concentrations of AVP for 24 h. (n=3). D) Fold change *Fgf23* levels in UMR-106 cells treated without (Ctrl) or with 100 nM AVP for indicated durations (n=3). E) Fold change *Fgf23* levels in UMR-106 cells treated without (Ctrl) or with the indicated concentrations of V_1a_ gonist, ([Phe^2^]OVT, [Phe^2^,Orn^8^]vasotocin) for 24 h. (n=3). F) Fold change *Fgf23* levels in UMR-106 cells treated without (Ctrl) or with the indicated concentrations of V_2_ agonist, ddAVP for 24 h (n=3). G) Serum cFGF23 and iFGF23 levels in wild-type and V_1a_ KO mice (n=4-5, each group).

### Tonicity-mediated FGF23 regulation is NFAT5-dependent

Our results show that elevated [Na^+^] levels (+20 mM) suppressed FGF23 formation, whereas low [Na+] levels (−20 mM) led to an increase in FGF23 synthesis. These bidirectional changes in FGF23 were inversely proportional to NFAT5 activity. In mice, *Fgf23* is expressed only in limited tissues such as the calvaria, spleen, and thymus, whereas *Nfat5* is almost ubiquitously expressed, including in the organs that produce *Fgf23* (Fig. S1). Therefore, NFAT5 may regulate FGF23 production. To test whether [Na^+^] mediated FGF23 changes were mediated by NFAT5 or not, we used the Crispr-Cas9 technology to knockout NFAT5 in UMR-106 cells *(NFAT5^KO^).* A single-cell derived clone of *NFAT5^KO^* was generated with a complete knockout, confirmed by immunoblotting (Fig. 4A) and qRT-PCR (Fig. 4B). Unlike in control cells, +NaCl treatment did not affect NFAT5 protein or mRNA expressions in *NFAT5^KO^* cells (Fig 4A/B). Moreover, cellular damage measured by LDH activity was more prominent in *NFAT5^KO^*cells than control cells, when treated with very high NaCl levels. (Fig 4C). These observations confirmed and validated successful NFAT5 deletion in UMR-106 cells. Importantly, *Fgf23* mRNA measured by qRT-PCR showed blunted response to tonicity in *NFAT5^KO^* cells compared with control cells (Fig. S2). However, the tonicity response to *Fgf23* was not completely abolished in *NFAT5^KO^*cells.

**Fig. 4.**
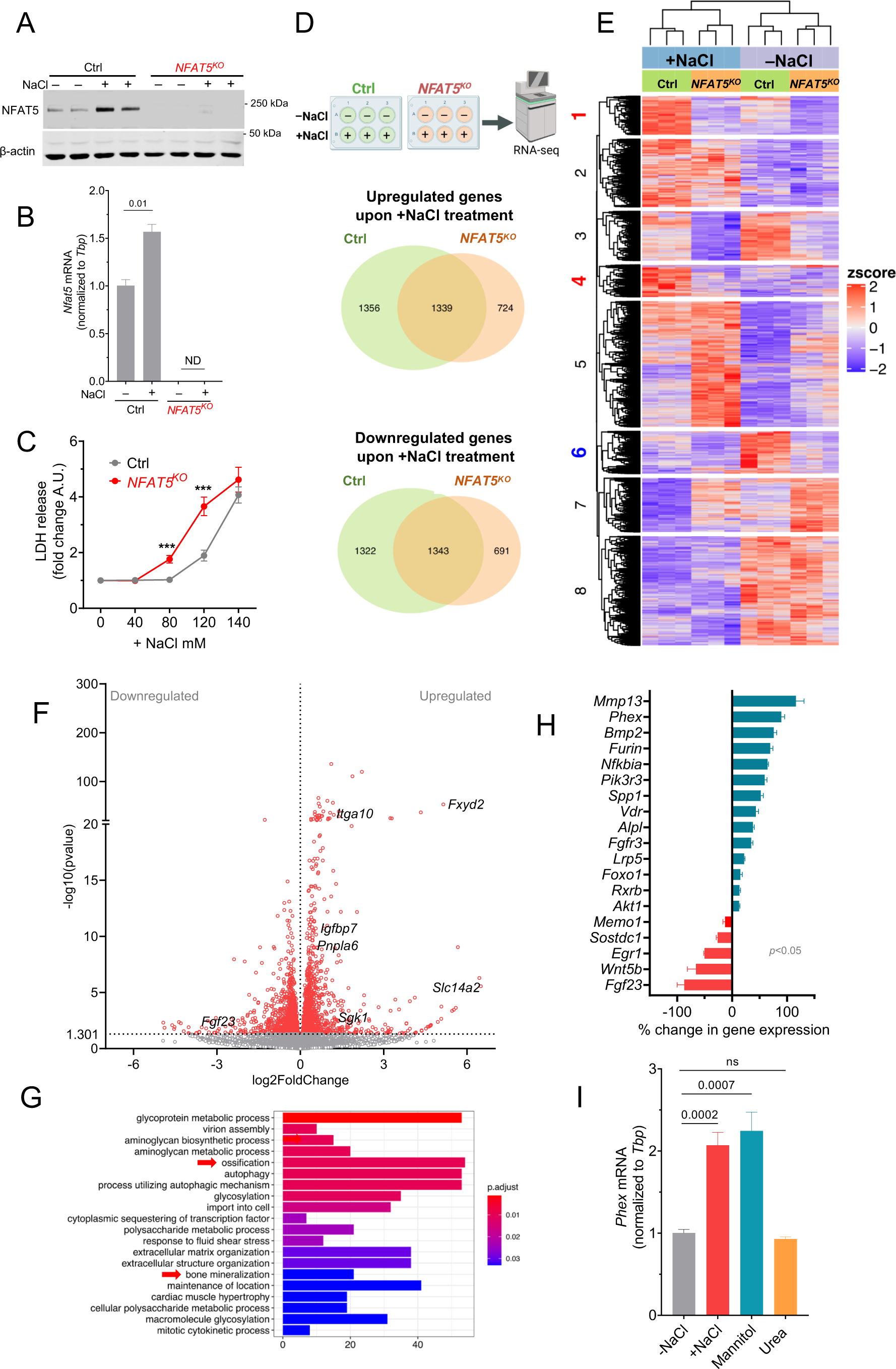
The [Na^+^] mediated regulation of FGF23 in UMR-106 cells requires NFAT5. A) Original immunoblot of NFAT5 and β-actin after -NaCl (−20 mM) and +NaCl (+20 mM) treatment for 24 h in control and *NFAT5^KO^* cells. B) *Nfat5* mRNA levels in control and *NFAT5^KO^* cells after -NaCl and +NaCl treatment for 24 h (n=3). *Nfat5* mRNA levels in *NFAT5^KO^* cells were not detected. C) LDH release in cell supernatants treated with different NaCl concentrations (n=3). ***(*p* < 0.001) indicate statistically significant difference between LDH levels in control and *NFAT5^KO^* cell supernatant at that particular +NaCl concentration. D) Venn diagrams showing a number of upregulated/downregulated genes in -NaCl vs. +NaCl treatment for 24 h. E) Heatmap of gene expression levels in control and *NFAT5^KO^* cells upon -NaCl vs. +NaCl treatment. F) Volcano plot of upregulated/downregulated genes in control cells upon -NaCl vs. +NaCl treatment. These genes were differentially expressed in control, but not in *NFAT5^KO^* cells upon -NaCl vs. +NaCl treatment. G) GO analysis of the upregulated genes in control cells that remain unchanged in *NFAT5^KO^*cells upon - NaCl vs. +NaCl treatment. H) FGF23-regulating genes that are statistically significantly (*p* < 0.05) upregulated/downregulated in control cells but remain unchanged in *NFAT5^KO^*cells upon -NaCl vs. +NaCl treatment. I) *Phex* mRNA levels measured by qRT-PCR upon -NaCl vs. +NaCl for 24 h in UMR-106 cells. Osmolality was corrected to +NaCl by adding mannitol or urea (n=3).

Next, we employed RNA-Seq to generate a comprehensive, unbiased survey of genes regulated by low vs. high [Na^+^] in UMR-106 cells in control and *NFAT5^KO^* cells. The RNA-seq of control vs. *NFAT5^KO^* cells revealed differential gene regulation upon -NaCl vs. +NaCl treatment as depicted in Venn diagrams (Fig. 4D). This suggested that transcriptionally active NFAT5 alters the expression of numerous genes in response to +NaCl (SI dataset 1/2). In the same line, in response to +NaCl treatment, control cells exhibited significant changes in a total of 5360 genes (*p*<0.05), whereas *NFAT5^KO^* cells showed alterations in 4097 genes (Fig. S3). Block 1/4/6 of the heatmap (Fig. 4E) suggested that genes that are either up or downregulated upon +NaCl treatment in control cells but mostly unchanged in *NFAT5^KO^* cells. Therefore, we focused only on these unique sets of potential NFAT5 targets (SI dataset 3). These NFAT5-target genes (n=2615) are further summarized in a volcano plot (Fig. 4F). *Slc14a2, Fxyd2, Sgk1, Pnpla6, Igfbp7,* and *Itga10* are known NFAT5 targets^37^, and our study confirmed this. Importantly, only control cells showed statistically significant downregulation of *Fgf23* after +NaCl treatment. The gene ontology (GO) analysis of the upregulated potential NFAT5 targets indicated the activation of ‘ossification’ and ‘bone-mineralization’ signaling pathways (Fig. 4G, SI dataset 4). After manually screening the genes involved in these pathways, we found that many of the identified genes regulate FGF23 and/or bone mineralization (Fig. 4H). Genes like *Phex, Mmp3, Nfkbia, Foxo1,* and *Akt1* are known to regulate FGF23 formation^38^. As proof of concept, we analyzed *Phex*, a known suppressor of FGF23, and found that *Phex* mRNA is significantly upregulated by the treatment of +NaCl and mannitol but not by urea (Fig. 2G, Fig. S6), confirming it’s a tonicity or NFAT5 target. However, other known regulators of FGF23 such as *Ibsp, Enpp1, Ankh, Pth1r, Dmp1, Fgfr1*, and *Sost*, were altered in both control and *NFAT5^KO^* cells (Fig. S4), suggesting NFAT5-independent mechanism in FGF23 regulation by [Na^+^]. Fig. S5 summarized potential NFAT5-target genes, which were previously postulated in [Na^+^] sensing and [Na^+^] homeostasis. Fig. S7 and S8 illustrate the analysis of all the KEGG and GO pathways, respectively.

### Hyponatremia patients have higher FGF23 levels

To test for the potential relevance of our findings *in vitro* for the situation *in vivo*, we investigated in a pilot study the levels of FGF23 in hyponatremic patients (with GFR>60) and compared them to the values of healthy subjects. The description of matching healthy controls and patients, including the etiology of hyponatremia, is provided in Table S3 and S4 respectively. As expected, serum [Na^+^] levels in hyponatremic patients were significantly lower than in the control group (Fig. 5A). Serum cFGF23 and iFGF23 isoforms were measured by ELISAs. As shown in Figure 5B, hyponatremic patients exhibited a significant increase in serum cFGF23 levels. Although there was a trend of increased serum iFGF23 in hyponatremic patients, the difference was not statistically significant (Fig. 5C). To evaluate the impact of altered FGF23 levels on phosphate homeostasis in these patients, we measured serum phosphate (Pi) levels. However, as shown in Fig. 5D, no significant difference was observed in serum phosphate levels between the control and hyponatremic patients. To perform correlation analysis, we plotted serum [Na^+^] against serum cFGF23 and iFGF23. Both cFGF23 and iFGF23 demonstrated a negative correlation with serum [Na^+^], but only cFGF23 exhibited a statistically significant correlation with [Na+] (Fig. 5E).

**Fig. 5.**
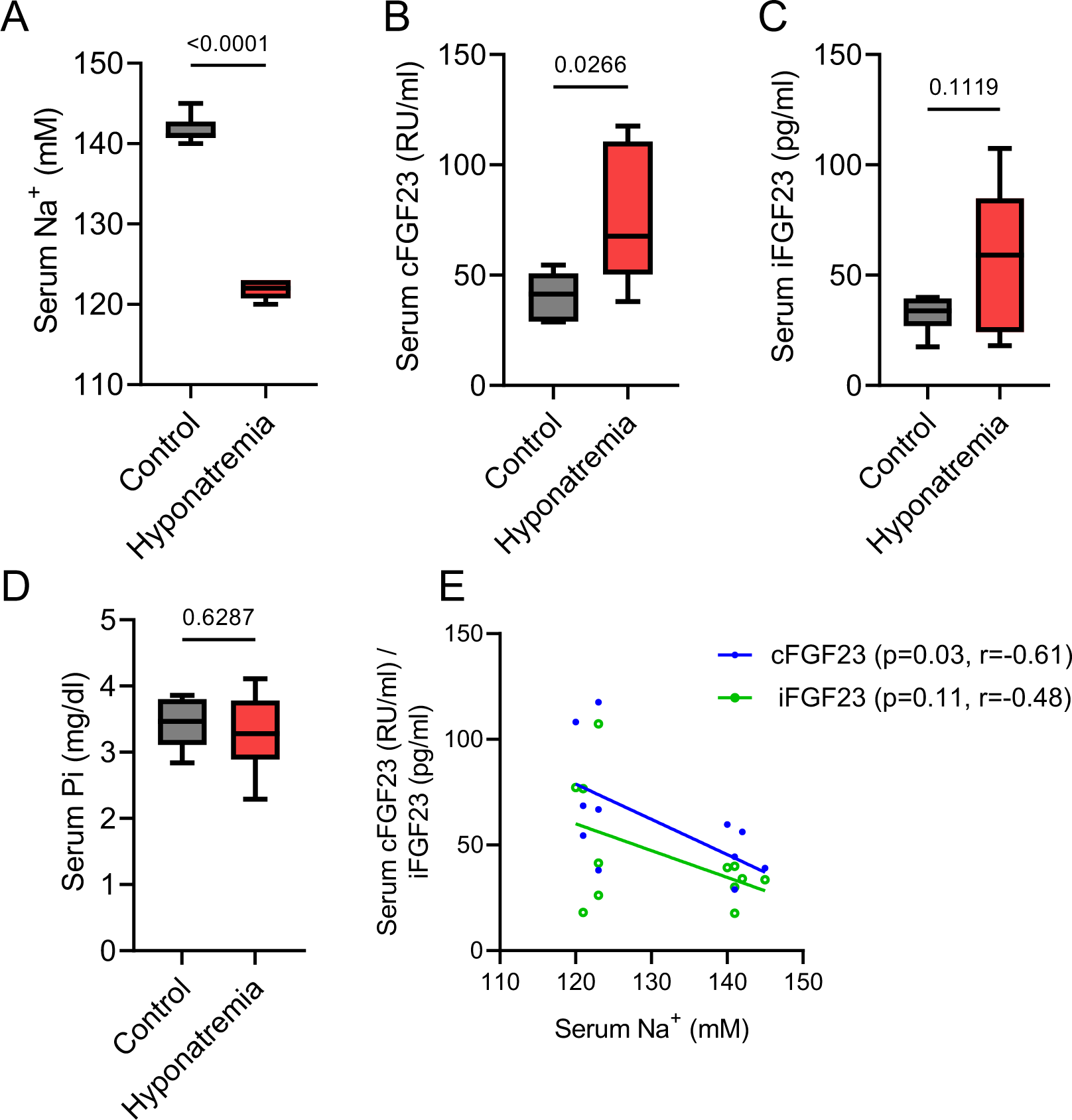
Hyponatremia patients have higher FGF23 levels. Serum [Na^+^] (A), cFGF23 (B), iFGF23 (C), and phosphate (D) in healthy matching controls and hyponatremic patients (n=6, each group). E) Correlation analysis between serum [Na^+^] and serum cFGF23/iFGF23. r= −0.6138, *p* value (two-tailed)= 0.0337 for [Na^+^] vs cFGF23; r= −0.4830, *p* value (two-tailed)= 0.1117 for [Na^+^] vs iFGF23.

## DISCUSSION

Our experiments demonstrate that physiologically and pathophysiologically relevant changes in [Na^+^] directly regulate FGF23 formation by osteoblasts. By manipulating the culture media using +NaCl, mannitol, or urea we showed that the cellular adaptation to tonicity, rather than osmolality, plays a significant role in regulating FGF23 levels upon change in [Na^+^]. Additionally, comprehensive RNA-seq analysis and NFAT5 knockdown experiments defined the role of the osmo-sensitive transcription factor NFAT5 in FGF23 regulation. We found no evidence that AVP, which increases in hyponatremia, regulates FGF23 production in the osteoblast cell line UMR-106.

We found that FGF23 regulation by tonicity is bidirectional, i.e. high [Na^+^] suppresses FGF23 while low [Na^+^] elevates FGF23 production, which is dependent upon NFAT5 activity. Under isotonic conditions, NFAT5 is both cytosolic and nuclear^32, 39^. With an increase in tonicity, it translocates to the nucleus, initiating the activation of a group of genes responsible for protecting cells against osmotic stress. This activation occurs through NFAT5 binding to a specific regulatory element called the osmotic response element or the tonicity-responsive enhancer, which is located in the promoter region of these genes^40–42^. As a result, a tonicity-dependent increase in nuclear NFAT5 is proposed as the key event in driving its target genes. While hypertonic stress promotes nuclear translocation and activates enhancer activity of NFAT5, hypotonicity induces its nuclear export supporting bidirectional regulation in response to tonicity changes^32, 39, 43^. The c-terminus of NFAT5 also contains a transactivation domain (TAD) and its activity varies directly with [Na^+^] concentration (low to high)^43^. NFAT5 regulates numerous gene expressions through its nuclear translocation during hypertonicity and nucleocytoplasmic export during hypotonicity. The +NaCl treatment in UMR-106 cells likely triggered a transcriptomics response via NFAT5, potentially impacting FGF23 regulation. Our study identified previously known genes that regulate FGF23 and bone mineralization as potential novel targets of NFAT5. For instance, *Phex* was one of the earliest genes identified to inhibit FGF23 formation^2, 3^. Now, we found that *Phex* is a NFAT5 target and gets remarkably upregulated by tonicity. Although our results demonstrated that FGF23 formation depends upon tonicity and NFAT5, the exact mechanism responsible for this phenomenon is unclear. Moreover, the tonicity response to *Fgf23* was not completely abolished in *NFAT5^KO^* cells, suggesting that NFAT5-independent mechanisms do also contribute to the regulation of *Fgf23* by [Na^+^].

Osmoregulation and [Na^+^] sensing mechanisms were mainly studied for cells in the hypothalamus (reviewed in^44^). In line with studies from others^27, 28, 30^, we found that osteoblasts can sense osmotically relevant changes in [Na^+^] and respond with an altered expression of several genes including *Fgf23*. Studies in RAW264.7 pre-osteoclastic cells showed increased osteoclast formation and resorptive activity in response to low [Na^+^] concentration^18, 19^. The molecular mechanism by which [Na^+^] is detected by bone cells is unknown. The expression of the osmoprotective transcription factor NFAT5 is regulated by local [Na^+^] content in many organs^45, 46^. Here, we demonstrate that the expression of this transcription factor in UMR-106 osteoblast-like cells is regulated by [Na^+^]. Previous studies focused on the effect of major changes in [Na^+^] (e.g. 50-100 mM NaCl hypo- and hypertonic conditions) on NFAT5 regulation^31, 32, 39^. However, such drastic changes of [Na^+^] occur only in the renal medulla, but not in bone. Now, our findings suggest that even differences of 20-40 mM NaCl significantly regulate NFAT5 activity in osteoblasts and osteoclasts, which is consistent with recent other studies^19, 30^. Serum/glucocorticoid regulated kinase 1 (*Sgk1*)^47^, transient receptor potential vanilloid type 4 (*Trpv4*)^48^, Na^+^/K^+^/ATPase (*Atp1a1*) and its associated protein such as sucrose non-fermenting-1-related serine/threonine kinase-SIK1 (*Snrk)*^49^ were previously postulated to take part in [Na^+^] sensing in various cells. Secondly, Na^+^/H^+^ exchanger activity^50^ (*Slc9a1*), and Na^+^/Ca^2+^ exchanger activity^51^ (*Slc8a1*) have been implicated in [Na^+^] homeostasis in bone cells. Our RNA-seq data show that, except for TRPV4, all these genes were significantly upregulated in control cells but not in *NFAT5^KO^* cells upon +NaCl treatment, indicating that these are NFAT5 targets (Fig. S5). Based on our findings and considering the role of NFAT5 as a master regulator of tonicity, it is tempting to speculate that changes in NFAT5 activation may be one of the earliest events in osteoblast adaptation to [Na^+^] levels. Therefore, NFAT5 likely plays a crucial role in [Na^+^] sensing in osteoblasts.

Our findings revealed that AVP does not take part in FGF23 formation. Most patients with chronic hyponatremia exhibit elevated levels of AVP that are inconsistent with the osmotic balance, even when SIAD is not the underlying cause of hyponatremia^52, 53^ (reviewed in^54^). The research group of Zaidi conducted seminal studies that have established a primary role for AVP signaling in bone mass regulation. Mice injected with AVP exhibited reduced osteoblast formation and increased osteoclast formation via the V_1a_ receptor and ERK signaling^24, 25^ (reviewed in^55, 56^). Conversely, mice injected with the V_1a_ antagonist SR49059 or genetic V_1a_ deficiency showed enhanced bone mass^24^. In contrast, the V_2_ receptor did not play a significant role in bone^25^. In line with these studies, we observed a significant expression of V_1a_ receptors but only a minimal expression of V_2_ receptors in UMR-106 cells. However, there was no evidence that AVP, V_1a_ agonist and V_2_ agonist regulate FGF23 production in UMR-106 cells. Consistently, serum FGF23 levels in V_1a_ KO mice were unchanged when compared to wild-type mice. These findings support a direct effect of [Na^+^] on FGF23 formation in hyponatremia. For instance, Verbalis *et.al*.^17, 57^ employed the SIAD model, where chronic hyponatremia induced by ddAVP and water loading promoted water retention. When water loading was not performed, the group of animals did not develop hyponatremia, and their bone mass did not show a significant reduction compared with the hyponatremia group. Moreover, cultured cells subjected to low [Na^+^] concentration exhibit increased bone resorption pathways and elicited gene expression changes driving osteoclast differentiation and functions^18, 19^, implying a direct effect of low [Na^+^] in bone loss. Overall, it seems that increased AVP levels play a significant role in bone loss among certain SIAD patients, but it may not take part in regulating FGF23 production.

We found a positive association between elevated FGF23 levels and hyponatremia in a pilot study conducted in humans. However, it remains unclear whether the increased FGF23 levels in hyponatremic patients are primarily influenced by low [Na^+^] and/or by the underlying cause of hyponatremia. Previously it was shown that in multivariate linear regression analysis, hyponatremia is independently associated with FGF23 level in patients with chronic systolic heart failure^34^. This further suggests that FGF23 elevation in hyponatremic patients might be due to low [Na^+^] levels. A larger sample size with hyponatremic patients will be necessary to confirm and unravel the possible relationship between [Na^+^] and FGF23 levels in humans. In conclusion, our study identified a crucial signaling between [Na^+^], NFAT5, and FGF23 formation, warranting further investigation to confirm its clinical significance.

## MATERIAL AND METHODS

All the chemicals were purchased from Sigma Aldrich unless otherwise stated.

### Culture of UMR106 cells

A rat osteoblastic cell line UMR-106 was originally purchased from ATCC (CRL-1661) and cultured at 5% CO_2_ in a growth medium consisting of Dulbecco’s Modified Eagle’s Medium DMEM, low glucose, GlutaMAX^TM^ (gibco; Cat. #21885-025) supplemented with 10% heat-inactivated FBS (Amimed; Cat. #2-01F30-I) and 100 U/ml penicillin, and 100 µg/ml streptomycin (gibco; Cat. #15140122). Cells between passages 4 and 10 were used in this study. When the cells reached near-confluency on 6-well plates, the cell medium was replaced with the experimental medium (2 ml/well) consisting of DMEM containing 1 nM of 1,25-dihydroxyvitamin D3 (Tocris; Cat. #2551), as described previously^58^. The cells were further treated with or without +NaCl, mannitol, or urea for specified concentrations and durations. For low-NaCl treatments, custom-made NaCl-free cell culture media was ordered commercially (Biotechne, Cat. #CUST07ATLB). The media formulation was developed to ensure that all the other components of the media were exactly similar to DMEM (gibco; Cat. # 21885-025), with the sole exception of the NaCl content. Thereafter NaCl was added to the medium to prepare a hypotonic medium to study the effect of low [Na^+^] concentration. The osmolality of all the culture media were measured on a Vapro 5600 (Wescor, Logan, UT) vapor pressure osmometer^59^. AVP (Tocris Cat. #2935) and ddAVP (Tocris Cat. #3396) were freshly prepared by dissolving them in water and treated for a specified duration and concentration. The V_1a_ agonist ([Phe^2^]OVT, [Phe^2^,Orn^8^]vasotocin) was a kind gift from Prof. Maurice Manning, The University of Toledo, USA. Its pharmacological properties in rat bioassay are reported earlier in a review^36^ by Manning *et.al*.

For FGF23 measurement in cell media, cells were cultured in a 100 mm petri dish. After 24 h of treatment with +NaCl, mannitol, and urea, cell media was concentrated using Pierce™ Protein Concentrator PES, 10K MWCO (Thermo Fisher; Cat. #88527). The resultant concentrate (150 µl) was used to measure c-terminal FGF23 by ELISA (Quidel; Cat. #60-6300).

### Animal experiments

To check *Fgf23* and *Nfat5* mRNA expression, organs were harvested from C57BL/6J wildtype mice (male, 8-12 weeks old) according to Swiss law and were approved by the veterinary administration of the Canton of Zurich (Kantonales Veterinäramt), Switzerland. The V_1a_ receptor KO mice were generated as previously reported^60^. The experiments with V_1a_ KO mice were conducted at the animal husbandry facility of the Teikyo University of Science. Around 400 µl of blood was drawn from 8-9 weeks old WT and V_1a_ receptor KO male littermates. The intact-FGF23 and c-terminal FGF23 were measured by ELISAs (Quidel, Cat. #60-6800, Quidel, Cat. #60-6300 respectively) according to the manufacturer’s instructions.

### LDH cytotoxicity assay

UMR-106 cells (control and *NFAT5^KO^* cells) were plated in a 12-well plate and cultured for 24 hr. Next, different concentrations of +NaCl were added to the culture media for a further 24 h. LDH was measured on aliquots of the +NaCl-treated cellular supernatant using the CyQUANT™ LDH Cytotoxicity Assay Kit (Thermo Fisher; Cat. #C20300). LDH activity was measured by measuring absorbance at 490 nm. The LDH release in +NaCl-treated cells is expressed in arbitrary units normalized to untreated cells.

### Immunoblotting

Cells were homogenized in ice-cold RIPA lysis buffer (Cat. #89900). Then cell lysates were centrifuged for 10 min at 2000 g and supernatants were used for immunoblotting. Protein concentration was measured by Bradford assay (CooAssay Protein Dosage Reagent; Uptima, Cat. #UPF86421). Equal amounts of protein (30 μg) were loaded in Laemmli buffer (pH 6.8) on 8% polyacrylamide gels. Electrophoretically separated proteins were blotted to nitrocellulose membranes at 100 V for 2 hours. Next, membranes were blocked in Odyssey blocking buffer (LI-COR; Cat. # 927-70001) for 1 hour and incubated with the diluted NFAT5 primary antibody (Thermo Fisher; Cat. #PA1-023) and β-actin (Santa Cruz; Cat. #sc-47778X) overnight at 4°C. On the following day, secondary antibodies -goat-anti-rabbit IRDye 800 (LI-COR; Cat. # 926-32211), goat-anti-mouse IRDye 680 (LI-COR; Cat. # 926-68070) were added to the membranes in Casein Blocking solution in deionized water (1:10) and incubated for 1 hour at room temperature. Then membranes were repeatedly washed with phosphate-buffered saline containing 0.1% Tween 20. The fluorescent signal was visualized using Odyssey IR imaging system (LI-COR Biosciences). Optical densities were quantified with ImageJ S5 and were normalized to β-actin.

### qRT-PCR

The RNA was extracted from UMR-106 cells and mouse tissues using a Nucleospin RNA isolation kit (Macherry Nagel; Cat. #740955). Rat tissue RNA was isolated as described in our previous study^61^. The RNA was transcribed using a High-Capacity cDNA Reverse Transcription Kit (Thermo Fisher; Cat. #4374966) and subjected to qPCR with SybrGreen Master Mix (Roche; Cat. #4707516001) using rat primers (Table 1). The relative quantification of gene expression based on double-delta Ct (threshold cycle) analysis was performed after normalization to *Tbp* expression.

**Table 1:**
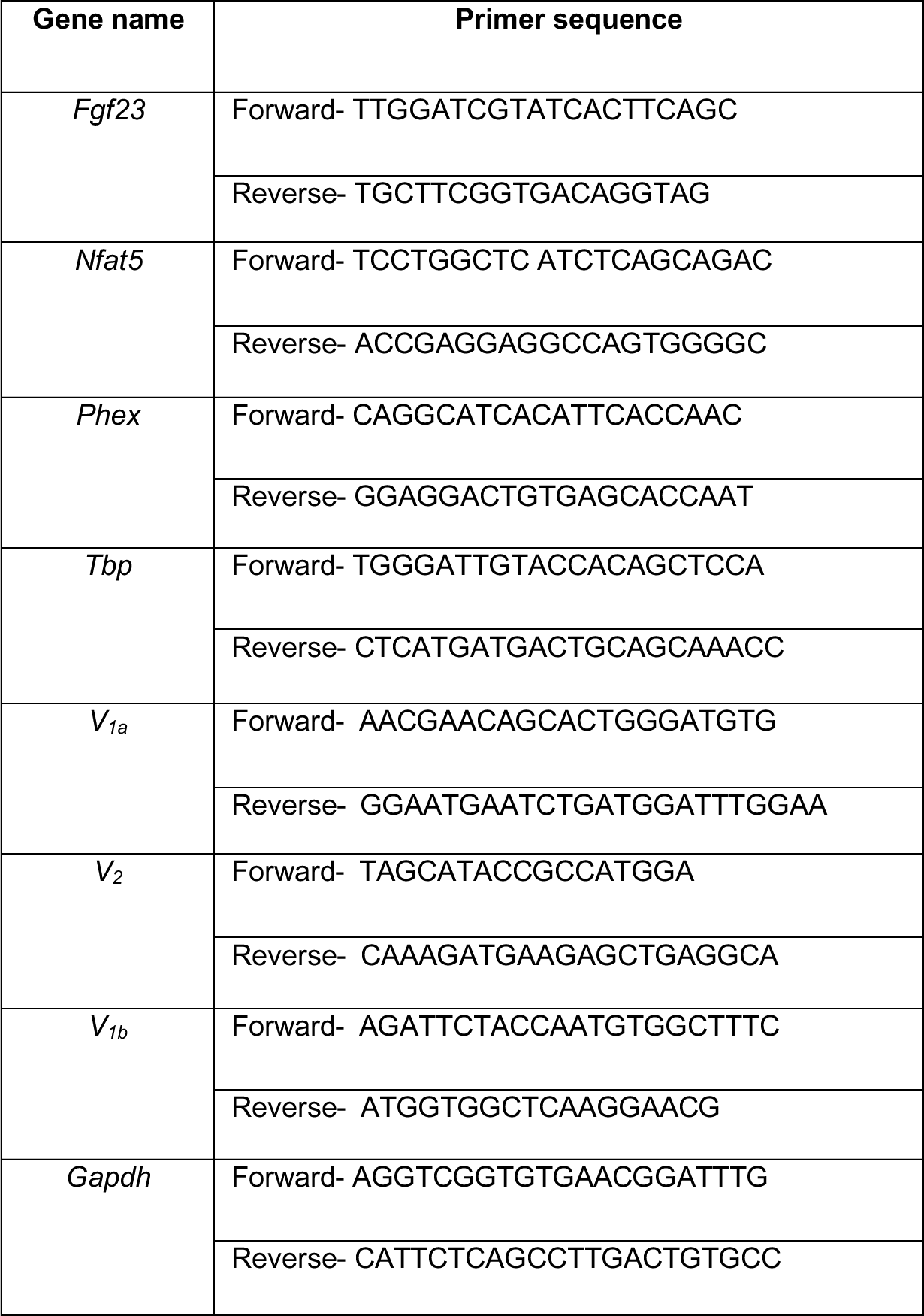
List of primers used for qPCR

### Generation NFAT5 knockout in UMR106 cells by CRISPR/Cas9

Single guide RNAs (sgRNAs) specifically targeting *Rattus norvegicus* NFAT5 at exon 1-4 were designed using the CRISPR designing tool (https://www.benchling.com/crispr). All sgRNA primers were synthesized by Microsynth AG, Balgach, Switzerland. sgRNA primers were (Table 2) cloned into BbsI-HF linearized pU6-(BbsI)_CBh-Cas9-T2A-mCherry (Addgene, Plasmid Cat. #64324) or pX330-U6-Chimeric_BB-CBh-hSpCas9 (Addgene, Plasmid Cat. #42230). Sanger sequencing (Microsynth AG) was used to verify the correct cloning of shRNA into the vectors. UMR106 cells were plated at 4 × 10^5^ per well in a 6-well plate and transfected with 2 µg of NFAT5 gRNAs per well using the FuGENE^®^ HD Transfection Reagent (Promega, USA). 24 h post-transfection, cells were trypsinized and single cells were sorted into a 96-well plate by FACS (BD BioSciences). A total of five clones derived from single-cell were propagated. NFAT5 knockout was confirmed by immunoblotting and qPCR. Following primers were used to confirm *Nfat5* deletion in *NFAT5^KO^* cells by qPCR: *Nfat5-KO* Fwd: GCCCTCGGACTTCATCTCAT; *Nfat5-KO* Rev: ACAGATTCTTCCAATAGTCCAGC

**Table 2:**
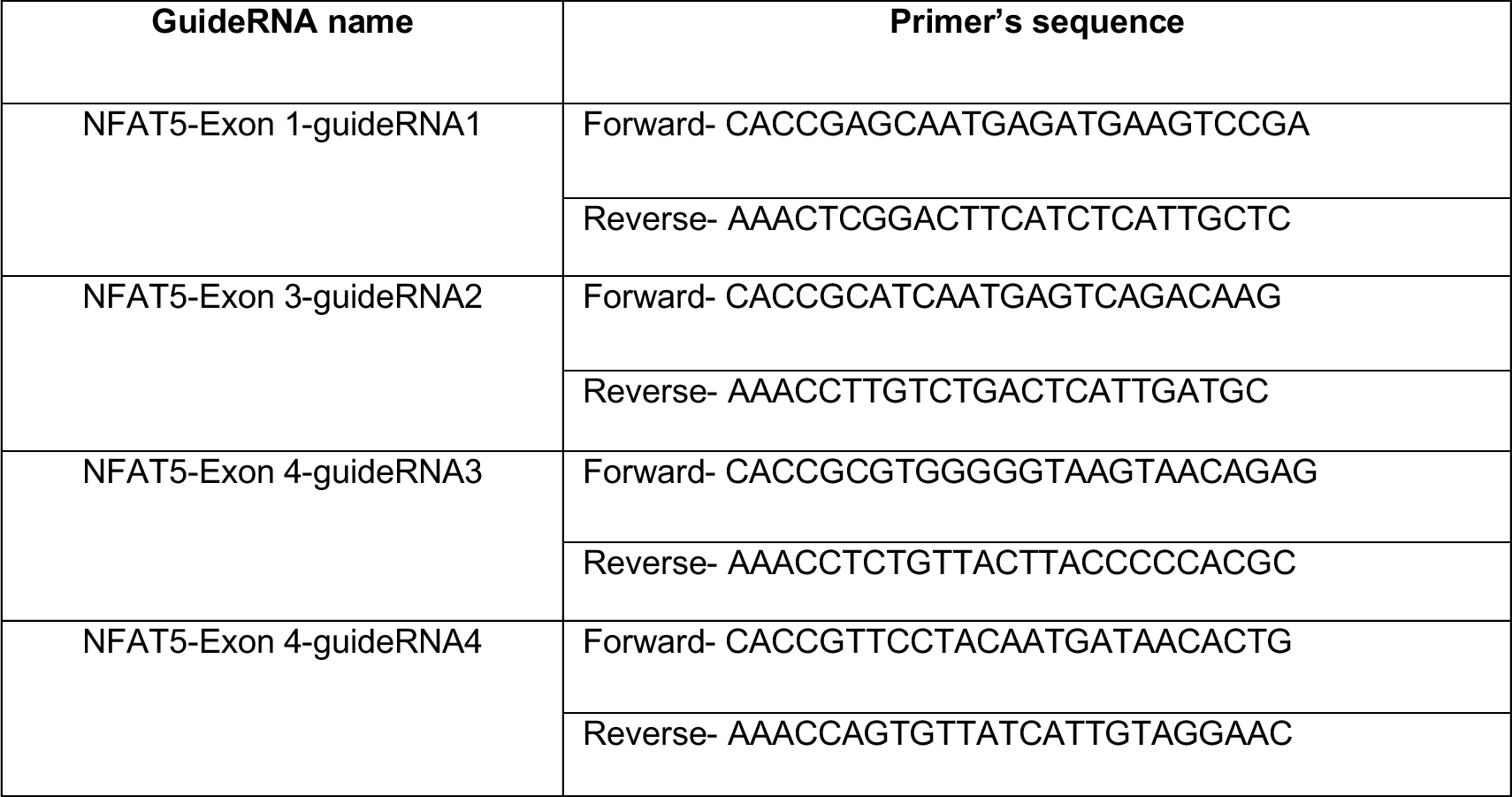
List of GuideRNA used for the generation of CRISPR/Cas9 knockout of NFAT5.

### RNA-sequencing and data analysis

Control and *NFAT5^KO^* cells were cultured as explained earlier. Cells were treated with low-NaCl and high-NaCl (difference of 40 mM NaCl) for 24 h. RNA was isolated using a Nucleospin kit as explained earlier. RNA-seq was performed commercially by Novogene (UK) Company Limited. In brief, amplified cDNA samples were subjected to different quality control standards. RNA sample was used for library preparation using NEB Next® Ultra RNA Library Prep Kit for Illumina®. Indices were included to multiplex multiple samples. Briefly, mRNA was purified from total RNA using poly-T oligo-attached magnetic beads. After fragmentation, the first strand cDNA was synthesized using random hexamer primers followed by the second strand cDNA synthesis. The library was ready after end repair, A-tailing, adapter ligation, and size selection. After amplification and purification, the insert size of the library was validated on an Agilent 2100 and quantified using quantitative qPCR. Libraries were then sequenced on Illumina NovaSeq 6000 S4 flow cell with PE150 according to results from library quality control and expected data volume.

We included 12 datasets in the RNA-Seq data analysis: three biological replicates in each group (control with -NaCl, control with +NaCl, *NFAT5^KO^*with -NaCl, and *NFAT5^KO^* with +NaCl). The fold changes were calculated by dividing the arithmetic mean of the normalized read counts by the number of replicates. The statistical significance of differential gene expression in RNA-Seq data was determined with *p* value <0.05. The supporting datasets S1 and S2 contain all of the genes identified upon +NaCl treatment in control and *NFAT5^KO^* cells respectively. While supporting datasets S3 contains unique genes, which were either significantly upregulated or downregulated in control cells but unchanged in *NFAT5^KO^* cells upon +NaCl treatment. Venn diagrams for the sets of genes up- or down-regulated were made by intersecting the lists of gene names between control and *NFAT5^KO^* cells (R package VennDiagram version 1.71). The heatmap of DEGs was made with the R package ComplexHeatmap (version 2.4.3) using as input the full set of genes differentially expressed between the normal and +NaCl conditions (p value < 0.05) for both control and *NFAT5^KO^* cells. The input expression matrix was normalized by row (gene) by computing a z-score. Both experimental groups (columns) and gene expression profiles (rows) were clustered using the euclidean distance and the hierarchical clustering algorithm. Gene set groups were obtained by cutting the dendrogram in eight slices with the cutree default implementation of the Heatmap function. Functional over-representation analysis of gene ontology biological processes was performed with the R package clusterProfiler (version 3.16.1) using as input the genes uniquely Up-regulated in the control cells upon +NaCl treatment (p value < 0.05) and considering a significance q-value cutoff of 0.05.

### Clinical study

Samples of patients with hyponatremia were collected during a prospective multicentric observational study (the Co-MED study, NCT01456533)^62^ conducted at the University Hospital of Basel Switzerland, and the Medical University Clinic Aarau, Switzerland, from June 2011 to August 2013. Samples of healthy controls were collected during a prospective double-blind, placebo-controlled randomized crossover study (the DIVE Study, NCT02729766)^63^ conducted at the University Hospital Basel, Switzerland, from March to June 2016. Venous blood samples were collected in the morning hours, between 8 am and 10 am. The local ethics committee (Ethic Committee of north-west Switzerland (EKNZ)) approved the study protocols and written informed consent was obtained from all study participants, including informed consent to further use of biologic material. Patients and healthy controls were matched based on age and BMI (n=6 each group). Detailed characteristics of the healthy controls and hyponatremia patients can be found in table S3 and S4, respectively. The serum electrolyte analysis [Na+] was performed immediately after collection on a ABL800FLEX blood gas analyzer (Radiometer, Thalwil, Switzerland). The serum was separated, and samples were stored at −80 °C until use. Commercial ELISA kits were employed to measure cFGF23 (Quidel, Cat. #60-6100) and iFGF23 (Quidel, Cat. #60-6600) in the serum according to the manufacturer’s instructions. Serum phosphate was measured by QuantiChrom™ Phosphate Assay Kit (Bioassay systems Cat. #DIPI-500).

## Supporting information

SI_Dataset S1_Ctrl_NaCl

SI_Dataset S2_KO_NaCl

SI_Dataset S3_NFAT5 targets

SI_Dataset S4.GO genes

## STATISTICS

All the gene or protein expression values are expressed as arithmetic means ± SEM, where n is the biological replicates. An unpaired Student’s t-test was used for comparisons between two groups using GraphPad Prism. In cases where the *p* value is not mentioned, the following applies: ns (not significant) *p* > 0.05, *\*p* ≤ 0.05, \*\**p* < 0.01, and \*\*\**p* < 0.001.

## ACKNOWLEDGMENTS

This work is supported by the Swiss National Science Foundation through the National Center of Competence in Research NCCR Kidney.CH. [grant number N-403-03-55 (to Ganesh Pathare)]. The authors sincerely thank Klaudia Kopper and Monique Carrel for their excellent technical assistance. Additionally, we express our gratitude to Prof. Maurice Manning from The University of Toledo, USA, for generously providing the V_1a_ agonist used in this study.

## AUTHORS CONTRIBUTIONS

GP conceptualized the study; ZR, EJ and GP designed the research; ZR, EJ, PK, SM, JR, MCC and GP performed the research. ASB performed RNA-seq data curation; HH, YK, JJ, and IRA contributed new reagents/analytic tools; ZR, EJ, LS, CAW, MAH, HMK, JL and GP analyzed data; ZR, EJ, and GP wrote the paper; LS, CAW, MAH, HMK, JL reviewed and edited the manuscript; GP was responsible for supervision; and GP was responsible for funding acquisition.

## DATA AVAILABILITY

All the relevant data associated with the current study can be found in the main text or supplementary material. The raw RNA-seq data used in this publication have been deposited in NCBI’s Gene Expression Omnibus and are accessible through GEO Series accession number GSE235694 (https://www.ncbi.nlm.nih.gov/geo/query/acc.cgi?acc=GSE235694). All data are available from the authors upon request.

