## Supplemental information for "Extracellular sodium regulates fibroblast growth factor 23 (FGF23) formation"

[1]Institute of Anatomy, University of Zurich, Zurich, Switzerland; [2]Swiss National Centre of Competence in Research "Kidney Control of Homeostasis", Switzerland; [3]Department of Biological Sciences, Ulsan National Institute of Science and Technology, Ulsan, Republic of Korea; [4]Membrane Transport Discovery Lab, Department of Nephrology and Hypertension and Department of Biomedical Research, Inselspital, University of Bern, Bern, Switzerland; [5]Department of Endocrinology, Diabetology and Metabolism, University Hospital Basel, Basel, Switzerland; [6]Department of Clinical Research, University of Basel, Basel, Switzerland; [7]Department of Animal Sciences, Teikyo University of Science, Yamanashi, Japan; [8]Institute of Clinical Microbiology and Hygiene, University Hospital of Regensburg and University of Regensburg, Regensburg, Germany; [9]Institute for Medical Microbiology, Immunology, and Hygiene, and Center for Molecular Medicine Cologne (CMMC), University of Cologne, Cologne, Germany; [10]Institute of Physiology, University of Zurich, Zurich, Switzerland

[†]contributed equally to this work

**Keywords:** FGF23, extracellular-sodium, hyponatremia, NFAT5, bone and kidney

**This PDF file includes:**

Supplementary tables and figures

### Supplementary tables

**Table S1**

| Cell Media | Osmolality (mOsm/Kg) |
|---|---|
| **Ctrl** | **300.7 ± 2.0** |
| **+NaCl (+20 mM)** | **345.7 ± 2.7** |
| **Mannitol (40 mM)** | **342.0 ± 1.5** |
| **Urea (40 mM)** | **342.3 ± 2.6** |

**Table S1.** Osmolality **±** SEM of the cell culture media used to study the effect of high extracellular [$Na^+$] (n=3, each group)

**Table S2**

| Cell Media | Osmolality (mOsm/Kg) |
|---|---|
| **Ctrl** | **302.3 ± 3.7** |
| **-NaCl (-20 mM)** | **260.7 ± 1.5** |
| **-NaCl + Mannitol (40 mM)** | **302.7 ± 1.9** |
| **-NaCl + Urea (40 mM)** | **298.0 ± 1.7** |

**Table S2.** Osmolality **±** SEM of the cell culture media used to study the effect of low extracellular [$Na^+$] (n=3, each group)

**Table S3**

| Nr | Age | Sex | Weight (Kg) | Height (cm) | Serum $[Na^+]$ (mM) |
|---|---|---|---|---|---|
| **1** | 30 | m | 86 | 185 | 145 |
| **2** | 49 | f | 68.4 | 160 | 141 |
| **3** | 40 | m | 65.8 | 169 | 142 |
| **4** | 50 | m | 71.2 | 182 | 141 |
| **5** | 48 | m | 91.8 | 186 | 140 |
| **6** | 30 | m | 62.9 | 165 | 141 |

**Table S3.** Characteristics of the matching healthy control group.

**Table S4**

| Nr | Age (y) | Sex | Weight (Kg) | Height (cm) | Serum $[Na^+]$ (mM) | Copeptin (pmol/L) | Etiology of Hyponatremia |
|---|---|---|---|---|---|---|---|
| 1 | 33 | m | 81.7 | 182 | 120 | 3.2 | SIADH (cause unknown) |
| 2 | 54 | m | 60 | 178 | 121 | 149 | Hypertonic translocation hyponatremia due to hyperglycemia |
| 3 | 47 | m | 70 | 170 | 123 | 110 | Hypovolemic Hyponatremia due to gastrointestinal fluid loss |
| 4 | 58 | m | 64 | missing | 123 | 11.9 | Hypovolemic Hyponatremia due to gastrointestinal fluid loss and mineralocorticoid deficiency |
| 5 | 55 | f | 55.5 | 152 | 121 | missing | Polydipsia (High Water Low Solute) |
| 6 | 55 | f | 35 | 170 | 123 | 2 | Polydipsia (High Water Low Solute) |

**Table S4.** Characteristics of the hyponatremic patients group.

### Supplementary figures

#### Fig. S1

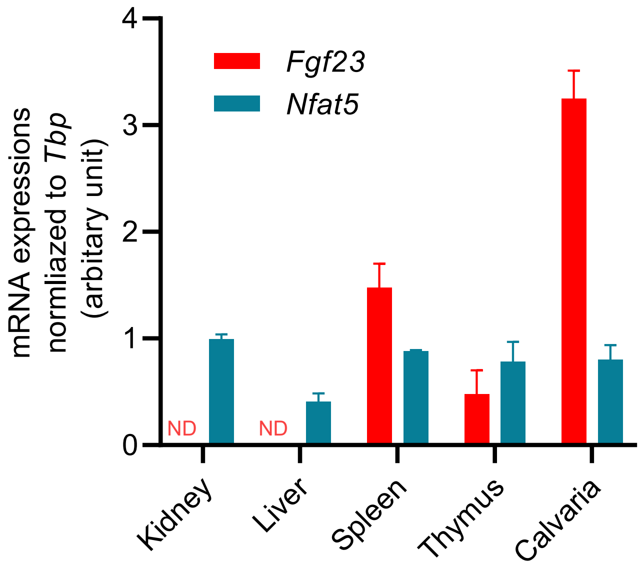


**Fig. S1.** The *Nfat5* and *Ffgf23* mRNA was measured by qRT-PCR in mouse kidneys, liver, spleen, thymus, and calvaria. Both *Nfat5* and *Ffgf23* mRNA were expressed in the spleen, thymus, and calvaria, while *Fgf23* mRNA was not detected (N.D) in the kidney and liver (n=3, each group).

#### Fig. S2

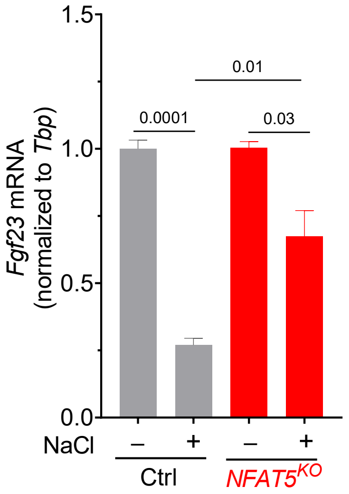


**Fig. S2.** *Fgf23* mRNA levels measured by qRT-PCR in control and *NFAT5*$^{KO}$ UMR-106 cells after -NaCl (-20 mM) vs. +NaCl treatment (+20 mM) for 24h (n=3, each group).

**Fig. S3**

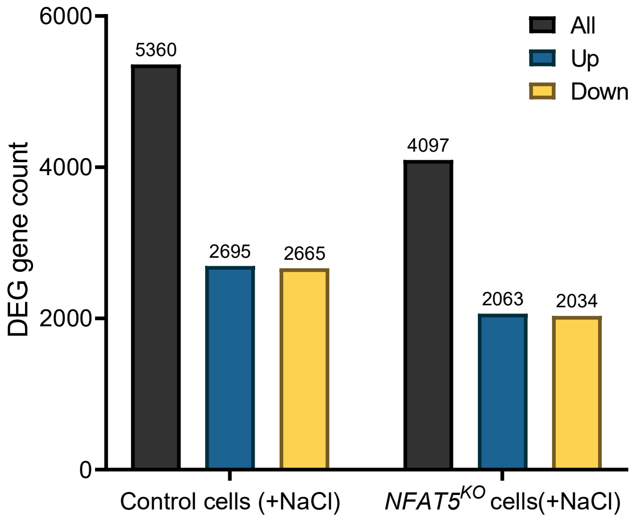


**Fig. S3.** Differentially expressed genes (DEG) count in control and *NFAT5*$^{KO}$ UMR-106 cells upon -NaCl vs. +NaCl treatment.

**Fig. S4**

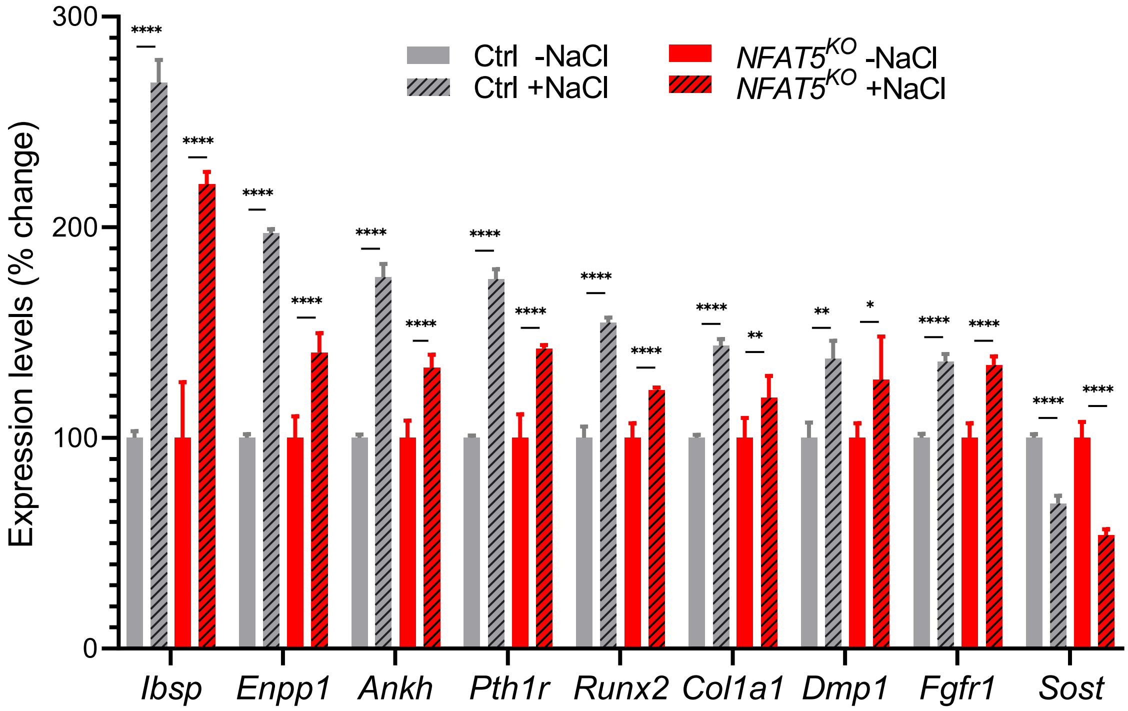


**Fig. S4.** The RNA-seq analysis resulted in DEG implicated in FGF23 regulation in both control and *NFAT5*$^{KO}$ UMR-106 cells. $p > 0.05$, $^{*}p \leq 0.05$, $^{**}p < 0.01$, $^{***}p < 0.001$ and $^{****}p < 0.0001$

**Fig. S5**

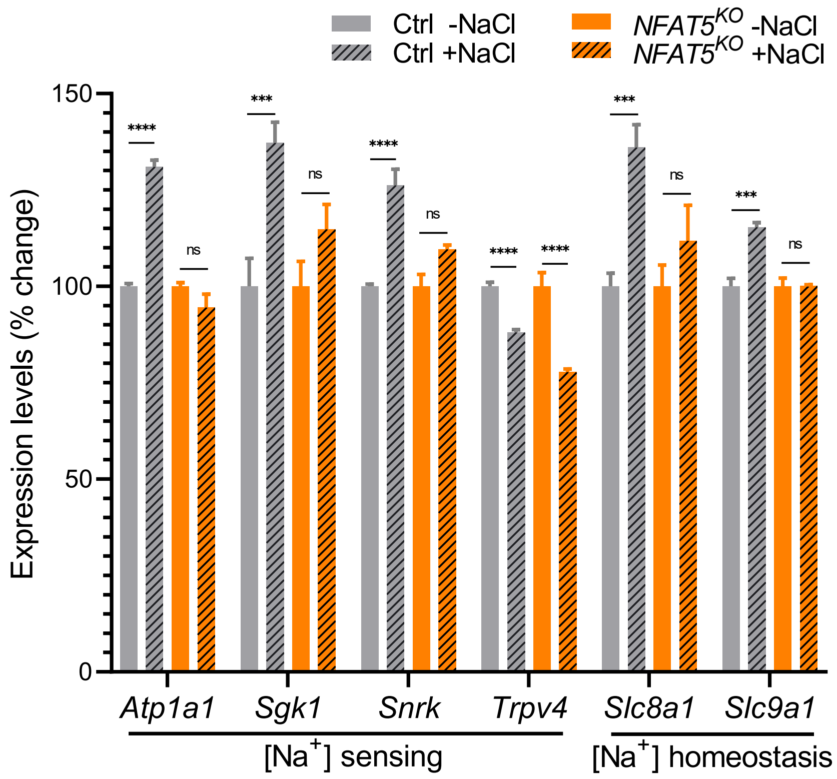


**Fig. S5.** The RNA-seq analysis resulted in DEG implicated in [$Na^+$] homeostasis in both control and *NFAT5*$^{KO}$ UMR-106 cells. (ns: not significant, $p > 0.05$, $*p \leq 0.05$, $**p < 0.01$, $***p < 0.001$ and $****p < 0.0001$)

**Fig. S6**

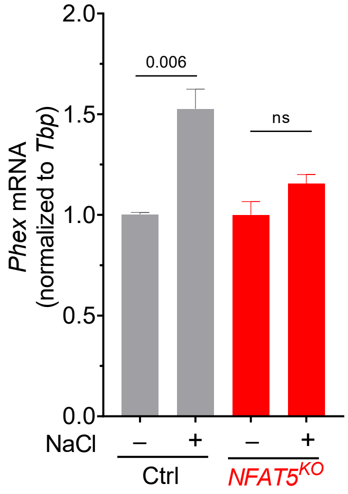


**Fig. S6.** *Phex* mRNA levels measured by qRT-PCR in control and *NFAT5*$^{KO}$ UMR-106 cells after -NaCl (-20 mM) vs. +NaCl treatment (+20 mM) for 24h (n=3, each group).

**Fig. S7**

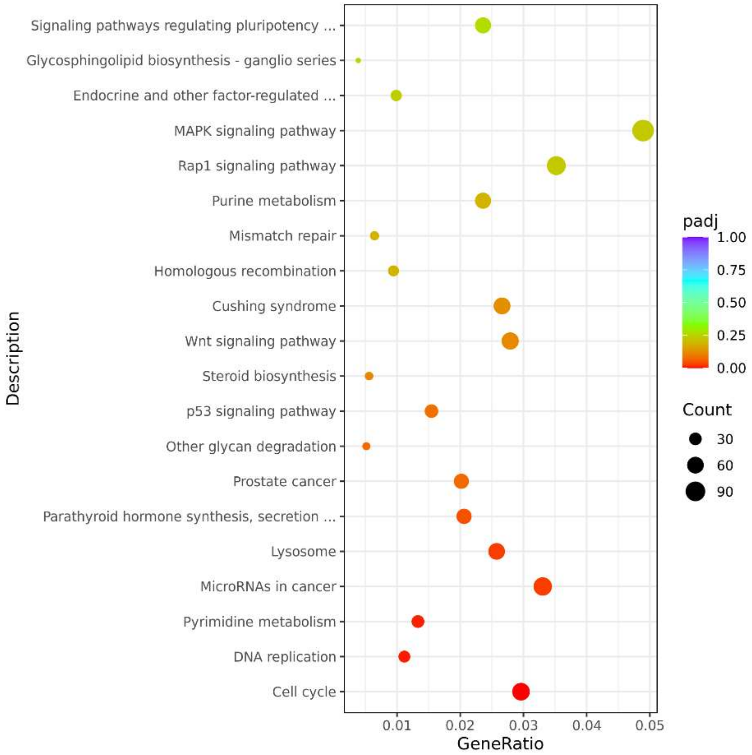


**Fig. S7. A)** KEGG pathways analysis in control UMR-106 cells upon +NaCl treatment.

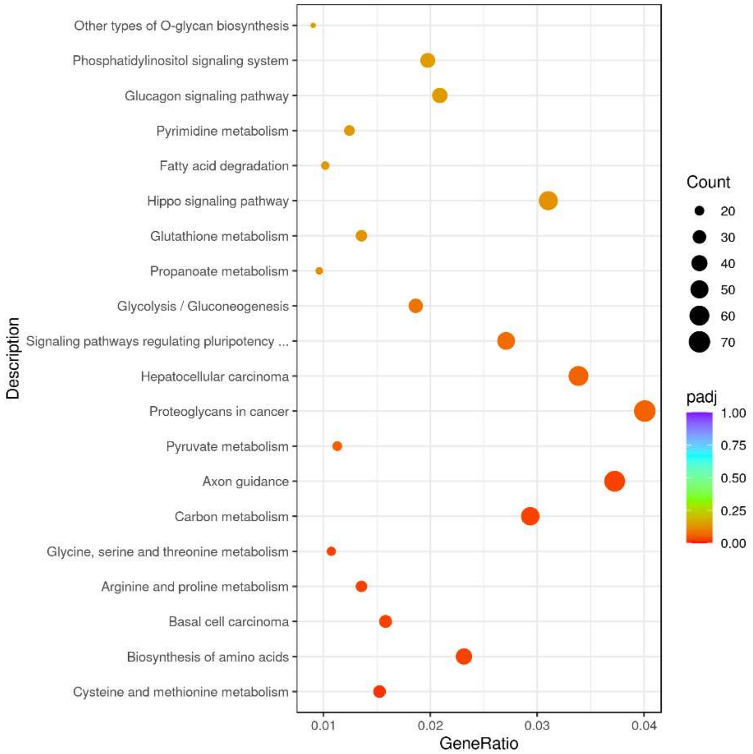


**Fig. S7. B)** KEGG pathways analysis in ***NFAT5**$^{KO}$* UMR-106 cells upon +NaCl treatment.

**Fig. S8**

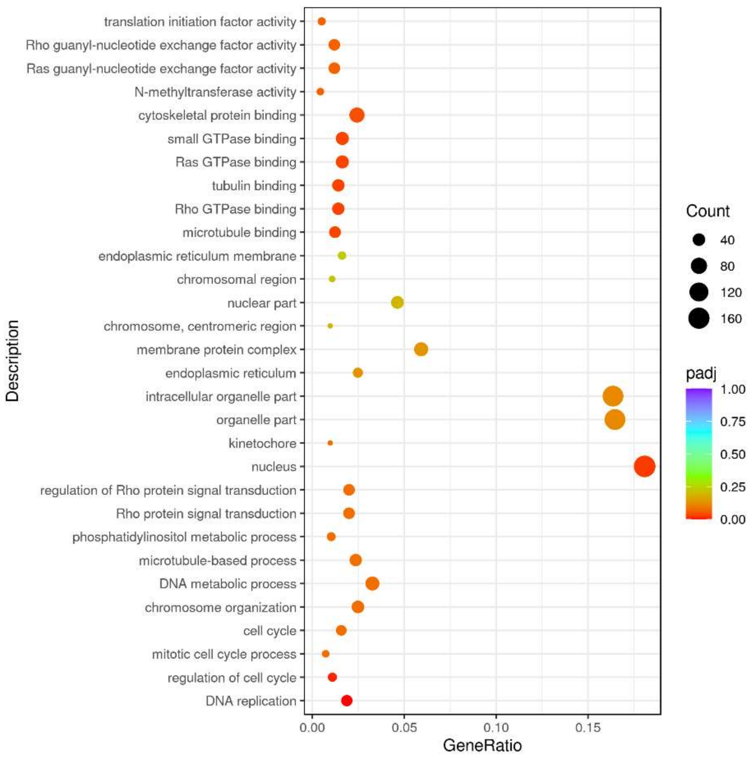


**Fig. S8. A)** GO pathways analysis in control UMR-106 cells upon +NaCl treatment.

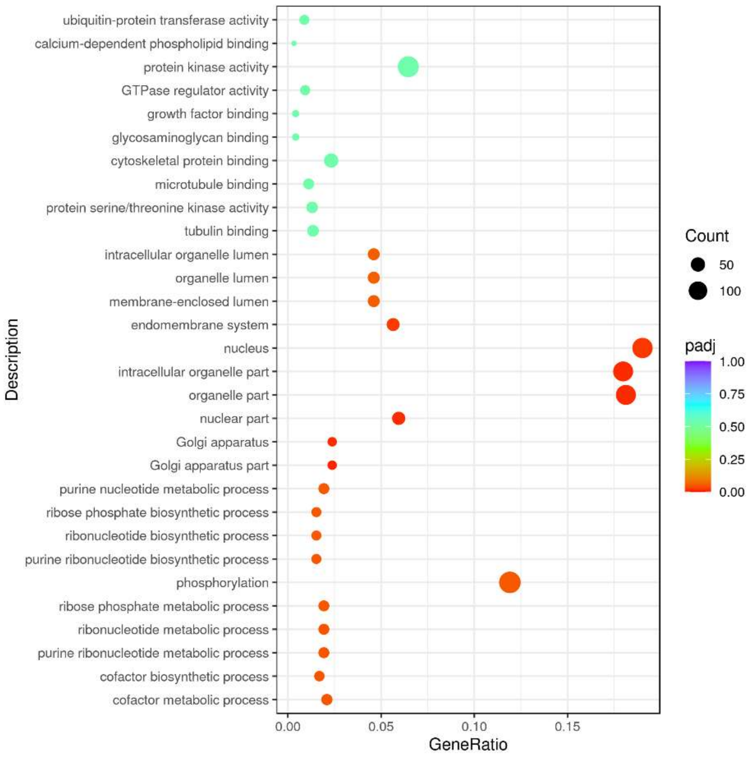


**Fig. S8. B)** GO pathways analysis in ***NFAT5$^{KO}$*** UMR-106 cells upon +NaCl treatment.
